# RNA splicing modulator induces peripheral neuropathy with increased neurofilament light chain levels via p53 signaling

**DOI:** 10.1101/2023.08.02.551640

**Authors:** Florian Krach, Tom Boerstler, Stephanie Reischl, Laura Krumm, Martin Regensburger, Jürgen Winkler, Beate Winner

## Abstract

RNA splicing modulators are a new class of small molecules with the potential to modify the expression levels of proteins. A recent clinical trial investigating the splicing modulator branaplam for Huntington’s disease to lower huntingtin levels was terminated due to the development of peripheral neuropathy. Here, we describe how branaplam leads to this adverse effect. On a cellular level, branaplam disrupts neurite integrity reflected by elevated neurofilament light chain levels in human induced pluripotent stem cell (iPSC)-derived motor neurons (iPSC-MN). Branaplam does not target neuropathy-associated genes. However, transcription factor binding site enrichment analysis indicates p53 activation. P53 activation upon branaplam treatment in iPSC-MN is linked to increased nucleolar stress, thereby enhanced expression of the neurotoxic p53-target gene BBC3. These findings illustrate that RNA splicing modulators may have clinically relevant off-target effects, implying the necessity of comprehensive pre-screening in human models prior to executing clinical trials.

**Graphical abstract:** 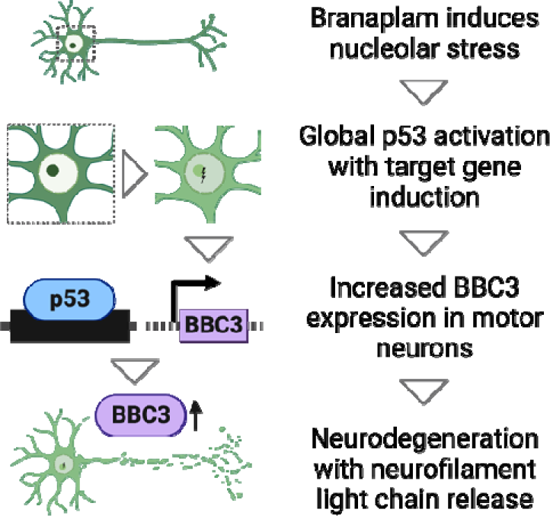

**One Sentence Summary:** Predicting side effects of RNA splicing modulator branaplam leading to neurotoxicity via nucleolar stress, p53 activation, and axonal degeneration.

## INTRODUCTION

Branaplam is a small molecule RNA splicing modulator and was discovered in a molecular splicing drug screen. It promotes changes in alternative splicing of the SMN2 mRNA ^1^ and underwent early clinical trials in SMA-affected infants (NCT02268552). Remarkably, one of branaplam’s off-target RNAs opened up an exciting chance to treat Huntington’s disease (HD). HD is a neurodegenerative disorder linked to CAG-repeat expansion in the huntingtin (HTT) gene, most likely leading to a toxic poly-Q protein. We and others have shown that branaplam induces a frame-shift-inducing pseudo-exon in the HTT transcript that leads to the destabilization of the HTT RNA and reduced mutant HTT protein expression ^2, 3^. Due to branaplam’s HTT-lowering activity, a phase II clinical trial was initiated for the treatment of HD (VIBRANT-HT; NCT05111249). On August 25^th^, 2022, the trial was suspended following the recommendation of the independent Data Monitoring Committee. In a very recent community update to the HD community, neurological symptoms and nerve conduction studies consistent with peripheral neuropathy were reported in conjunction with an increase in serum neurofilament light chain (NfL) in branaplam-treated individuals ^4^. However, it remains elusive how the RNA splicing modulator leads to peripheral neuropathy. To solve this issue, human-based models could be a crucial prerequisite in order to identify adverse events prior to initiating a clinical study.

Here, we describe the cellular mechanism of how the small molecule splicing modulator branaplam leads to peripheral neuropathy and NfL increase. Human iPSC-derived motor neurons treated with branaplam exhibit neurite fragmentation and increased NfL release into the medium. We re-analyze RNA-seq datasets of branaplam-treated human fibroblasts and identify a modest but clear signal of p53-target gene activation. We evaluate the origin of p53 activation and reveal increased nucleolar stress upon branaplam in human iPSC-derived motor neurons. Branaplam-treated neurons induce expression of the neurotoxic p53-target BCL2 binding component 3 (BBC3).

## RESULTS

To test whether cellular signs of peripheral axonopathy induced by branaplam can be measured in vitro, we differentiated induced pluripotent stem cells (iPSC) from healthy individuals into motor neurons (iPSC-MN) using a previously established protocol ^5^ and investigated the toxic effect on human neurites (Fig. 1a). In branaplam-treated iPSC-MN, we detected more disintegrated neurites, consistent with axonal degeneration observed in neuropathies (Fig. 1b and c). Additionally, levels of NfL in the supernatant of branaplam-treated iPSC-MN were quantified. Strikingly, we detected an increase in NfL levels in the supernatant (Fig. 1d). This illustrates that our cellular model recapitulates cytopathogenic features coherent with the clinically observed parameters of neuropathy and elevated NfL in the phase 2 HD clinical trial. To uncover a direct mRNA-mediated mechanism involved in this neuropathy, we leveraged our previously published human fibroblast and iPSC-neuron RNA-seq dataset of control (Ctrl) and HD patients treated with the low dose of 10 nM branaplam for 72 h resulting in a significant reduction of mHTT protein levels ^3^ (Supplementary Fig. 1a). First, we investigated if branaplam targets genes known to cause neuropathy in a monogenic manner by comparing our results with two independent genetic panels (Supplementary Fig. 1b). We found that one branaplam target also appeared in one of the neuropathy gene panels (Supplementary Fig. 1 c). Branaplam induces an in-frame exon inclusion in the Protein O-mannosyltransferase 2 (POMT2) (Supplementary Fig. 1d). The insertion does not lead to a frameshift or insertion of an in-frame premature STOP codon. While the functional consequence of this exon inclusion is not clear, POMT2 is linked to muscular dystrophies, i.e. dystroglycanopathies ^6, 7^. However, neuropathic features have not been described to our knowledge, making it an unlikely culprit. We further examined whether branaplam treatment affects the expression of neuropathy-associated genes in an indirect manner, but we did not observe substantial effects of branaplam (Supplementary Fig. 1e-h).

**Fig. 1.**
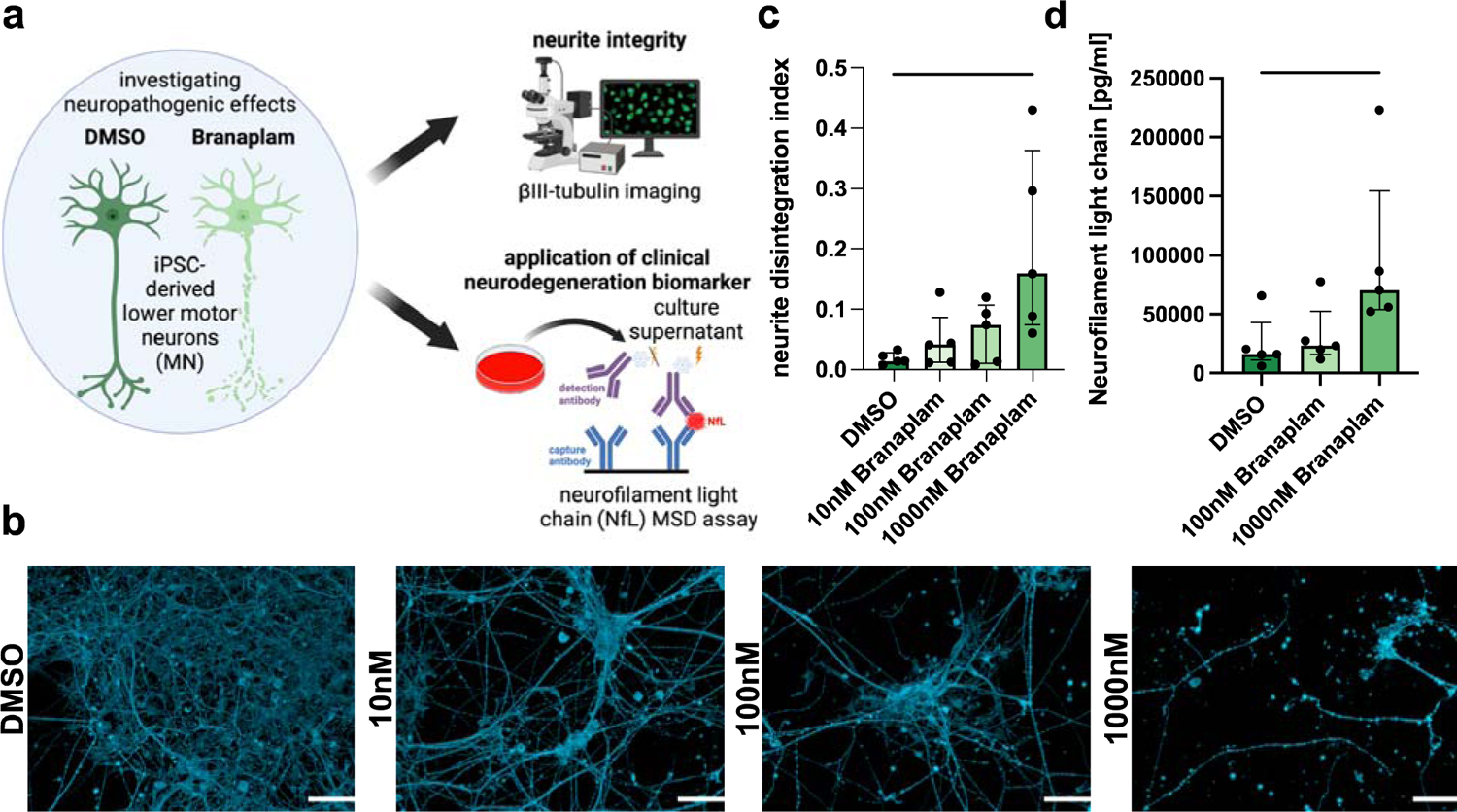
Branaplam leads to axonal degeneration and NfL increase in human iPSC-MN. (A) Paradigm illustrating experimental approach. (B) Representative pictures of beta-III-Tubulin in DMSO and 1000 nM-treated conditions. All shown images are from UKERiff2-X-18. Scale bar: 50 µm. (C) Bar plot showing quantification of neurite disintegration index (= disintegrated area (betaIII-Tubulin intensity above mean+3*standard deviation of DMSO) / total area (beta-III-Tubulin area). Dots depict individual values per cell line (n=5), graph as median with interquartile range. Statistics: Friedman test: P value: 0.0167. Dunn’s multiple comparison test: * < 0.05; ** < 0.01; *** < 0.001. (D) Bar plot depicting quantification of Neurofilament light chain (NfL) levels in media supernatant of iPSC-derived neurons after 5d of treatment with DMSO, 100 nM or 1000 nM branaplam. Statistics: Friedman test: P value: 0.0008. Dunn’s multiple comparison test: * < 0.05; ** < 0.01; *** < 0.001.

We reasoned that branaplam may mediate toxic neuropathic features via mechanisms other than targeting mRNA. Hence, we studied the cellular state of branaplam-treated fibroblasts by delineating the pathways altered upon treatment. Specifically, we investigated the transcription factor (TF) binding sites (TFBS) of differentially expressed genes (Fig. 2a-d). Genes that are downregulated upon branaplam treatment did not show a large number of significant enrichments of TFBS (Fig. 2d). However, genes that are upregulated upon branaplam exhibited a large number of highly significant TFBS (Fig. 2c). Strikingly, the tumor suppressor gene p53 was among the top 10 TFs with the highest percentage of shared genes. To confirm this global activation of p53, we integrated an ENCODE p53 ChIP-seq dataset ^8, 9^ and analyzed gene expression changes upon branaplam treatment in Ctrl and HD fibroblasts in genes with a p53 ChIP-seq peak in a location of up to 5000 bp upstream of the gene start. Strikingly, significantly higher gene expression was observed in Ctrl and HD fibroblasts upon branaplam treatment of p53 targets compared to a shuffled background (Fig. 2e and f). Besides the well-known transcriptional activation of CDKN1A (p21), p53-mediated gene activation is also known to upregulate BBC3, also known as PUMA. Interestingly, both CDKN1A and BBC3 were upregulated upon branaplam treatment in fibroblasts (Fig. 2g and h). Together these data suggest a global activation of p53 signaling. P53’s function as a tumor suppressor to attenuate proliferation is widely known. Hence, we predicted that fibroblasts treated with branaplam show an altered cell cycle. Since 10 nM branaplam did not yield proliferation changes in cortical progenitors ^3^, we performed the experiment using two higher doses (100 nM and 1000 nM) (Fig. 2i). To probe for shared mechanisms, we treated with paclitaxel, a cytostatic agent leading to an M-phase arrest by inhibiting the mitotic spindle apparatus, clinically used for the treatment of, e.g. ovarian and breast cancer. A common side effect is peripheral neuropathy. As expected, paclitaxel leads to an M-phase arrest. Branaplam also alters the cell cycle in a dose-dependent manner but leads rather to a G1/S-phase arrest. Together, this suggests that branaplam treatment induced cell cycle arrest via p53 activation but in a non-paclitaxel-like manner (Fig. 2i-l).

**Fig. 2.**
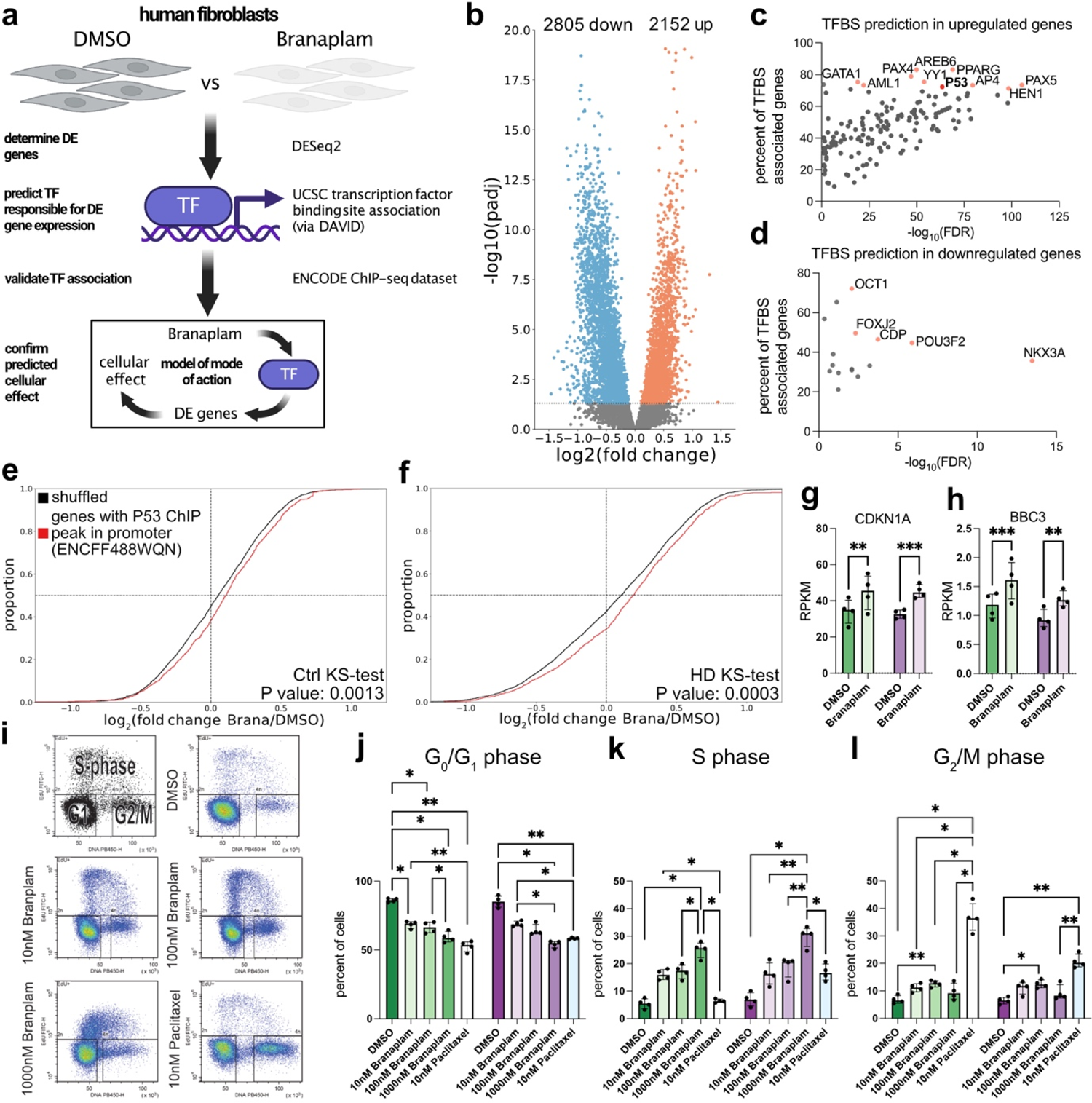
Branaplam leads to p53 activation and cell cycle arrest. (A) Experimental analysis strategy. DMSO vs branaplam treated fibroblast RNA-seq data ^3^ was used to determine differential gene expression and analysis of enrichment of transcription factor (TF) binding sites (TFBS), followed by validation using ENCODE ChIP-seq data and determine of the cellular mode of action (B) Volcano plot of differentially expressed genes (blue: downregulated, red: upregulated). Y-axis depicts negative log10 of the adjusted P value from the DESeq2 output. (C) Scatter plot of significantly enriched (x-axis, false discovery rate (FDR)) TFBS in genes upregulated upon branaplam treatment. Y-axis depicts percentage genes associated TF that are differentially upregulated upon branaplam. In red are 10 TFs with high significance and percentages. (D) Scatter plot of significantly enriched (x-axis, false discovery rate (FDR)) TFBS in genes downregulated upon branaplam treatment. Y-axis depicts percentage genes associated TF that are differentially downregulated upon branaplam. In red are 10 TFs with high significance and percentages. (E) Cumulative distribution plot of log2(fold changes) of branaplam vs. DMSO in Ctrls in genes with p53 ChIP-seq peak within region of 5000bp upstream of gene start (red, n=865 genes) or in a randomly shuffled background of equal size (black, n=865 genes). Significance calculated with 2 sample Kolmogorov-Smirnov (KS) test. (F) Cumulative distribution plot of log2(fold changes) of branaplam vs. DMSO in HD in genes with p53 ChIP-seq peak within region of 5000bp upstream of gene start (red, n=865 genes) or in a randomly shuffled background of equal size (black, n=865 genes). Significance calculated with 2 sample Kolmogorov-Smirnov (KS) test. (G) Bar plot of CDKN1A RPKM values with Ctrl (greens) and HD (purples) fibroblast with DMSO and branaplam (opaque) treatment. Dots depict individual values, graph as median with interquartile range. Statistics: paired 2-way ANOVA: P value (treatment): <0.0001; P value (disease): 0.8851; P value (interaction): 0.3535. Šídák’s multiple comparisons test: P value (Ctrl): 0.001; P value (HD): 0.0004. (H) Bar plot of CDKN1A RPKM values with Ctrl (greens) and HD (purples) fibroblast with DMSO and branaplam (opaque) treatment. Dots depict individual values, graph as median with interquartile range. Statistics: paired 2-way ANOVA: P value (treatment): <0.0001; P value (disease): 0.1239; P value (interaction): 0.3071. Šídák’s multiple comparisons test: P value (Ctrl): 0.0006; P value (HD): 0.002. (I) Paradigm of EdU FACS cell cycle analysis and example scatter plots and gatings from DMSO, 10 nM branaplam, 1000 nM branaplam and Paclitaxel of UKERf33Q. (J) Bar plot of percentages of cells in G0/G1 phase in Ctrl (greens) and HD (purples) treated for 72h with 10 nM, 100 nM, 1000 nM branaplam (opaques) or 24h 10 nM Paclitaxel (white/blue). Dots depict individual values, graph as median with interquartile range. Statistics: paired 2-way ANOVA: P value (treatment): <0.0001; P value (disease): 0.7840; P value (interaction): 0.0166. Šídák’s multiple comparisons test: * < 0.05; ** < 0.01; *** < 0.001. (K) Bar plot of percentages of cells in S phase in Ctrl (greens) and HD (purples) treated for 72h with 10 nM, 100 nM, 1000 nM branaplam (opaques) or 24h 10 nM Paclitaxel (white/blue). Dots depict individual values, graph as median with interquartile range. Statistics: paired 2-way ANOVA: P value (treatment): <0.0001; P value (disease): 0.0345; P value (interaction): 0.0018. Šídák’s multiple comparisons test: * < 0.05; ** < 0.01; *** < 0.001. (L) Bar plot of percentages of cells in G2/M phase in Ctrl (greens) and HD (purples) treated for 72h with 10 nM, 100 nM, 1000 nM branaplam (opaques) or 24h 10 nM Paclitaxel (white/blue). Dots depict individual values, graph as median with interquartile range. Statistics: paired 2-way ANOVA: P value (treatment): <0.0001; P value (disease): 0.0127; P value (interaction): <0.0001. Šídák’s multiple comparisons test: * < 0.05; ** < 0.01; *** < 0.001.

Multiple cellular dysregulations result in p53 activation. Since branaplam is an RNA-targeting molecule, we reasoned that an underlying RNA-associated mechanism is likely. In this respect, nucleolar stress is a prominent example. Here, dysregulation of rRNA transcription or metabolism in conjunction with other mechanisms disrupts the nucleolar structure and integrity, resulting in p53 activation. Nucleolar stress can be measured via the translocation of NPM1 from the nucleolus to the nucleoplasm ^10^. We sought to investigate whether this is occurring in human neurons upon branaplam exposure (Fig. 3a). iPSC-MN treated with 1000 nM of branaplam exhibited increased ratios of NPM1 signal in the nucleoplasm vs. the nucleolus (Fig. 3b and c). This indicates the induction of nucleolar stress in iPSC-MN. P53-mediated BBC3 expression is able to induce neurite degeneration in physiological and pathological conditions ^11–14^. We observed an increased BBC3 expression in the iPSC-MN treated with branaplam, indicative of activated p53 signaling (Fig. 3d and e).

**Fig. 3.**
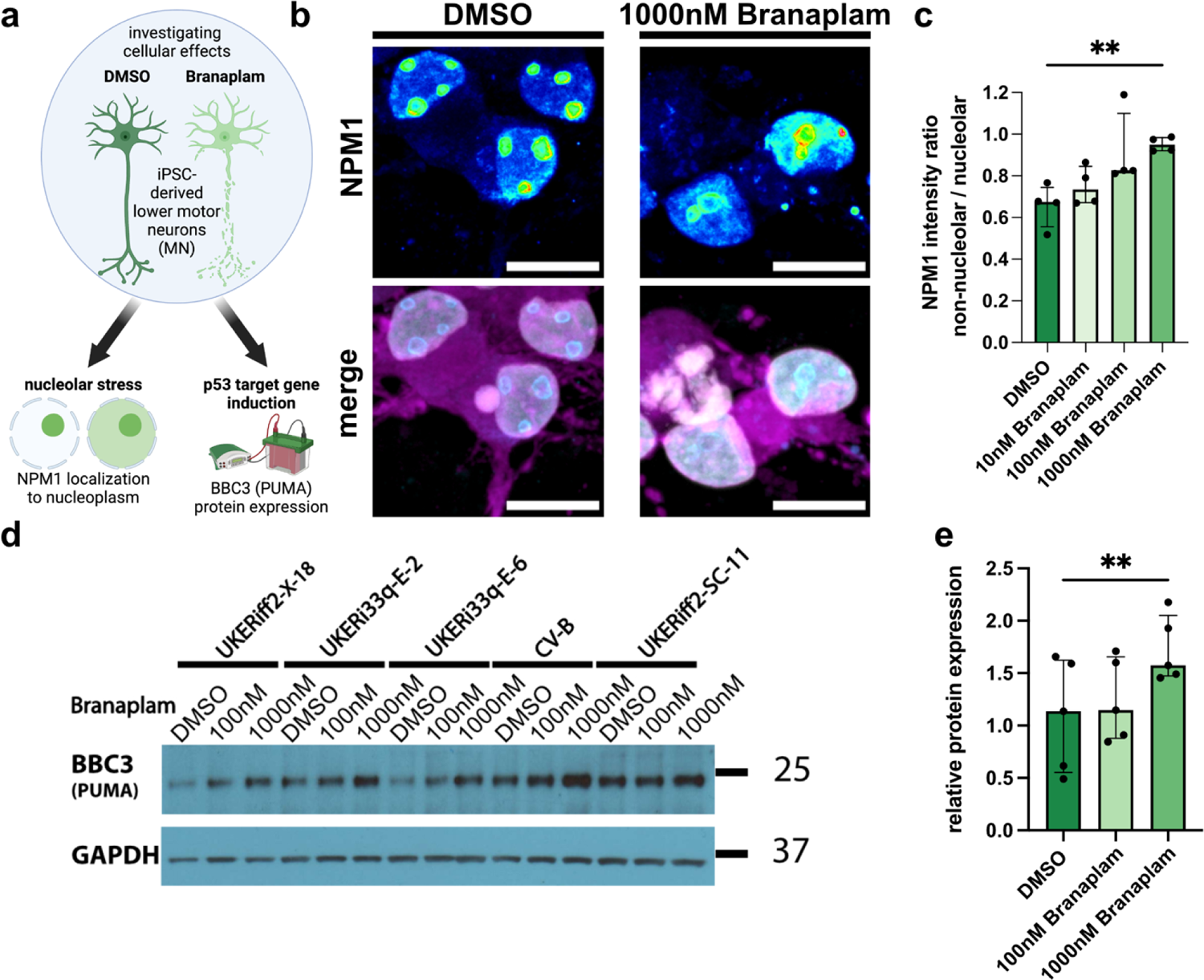
Branaplam leads to activation of nucleolar stress and expression of BBC3 and axonal degeneration in human iPSC-MN. (A) Paradigm illustrating experimental approach. (B) Representative pictures of betaIII-Tubulin-positive neurons (magenta) with NPM1 signal (LUT, blue=low fluorescence intensity, white=high fluorescence intensity) in the nucleus (white) of UKERiff2-X-18. Scale bar: 10 µm (C) Bar plot quantifying fluorescence intensity ratio of NPM1 in non-nucleolar nucleoplasm / nucleolar area (n=4 cell lines). Dots depict individual values per cell line, graph as median with interquartile range. Statistics: Friedman test: P value: 0.0062. Dunn’s multiple comparison test: * < 0.05; ** < 0.01; *** < 0.001. (D) Western blot of BBC3 (also known as PUMA) in iPSC-MN of 5 cell lines treated for 5 days with DMSO, 100 nM and 1000 nM branaplam. GAPDH serves as loading control. (E) Bar plot showing densitometric quantification of BBC3 signal normalized to GAPDH. Dots depict individual values per cell line, graph as median with interquartile range. Statistics: Friedman test: P value: 0.0008. Dunn’s multiple comparison test: * < 0.05; ** < 0.01; *** < 0.001.

## DISCUSSION

Our results indicate that this novel type of small molecule may have yet unknown off-targets that lead to clinically relevant side effects in adult patients, which was previously not noted. Branaplam has been first under clinical investigation in infants with SMA without severe adverse effects. However, due to three alternative options of causal SMA treatment, the development for SMA patients was discontinued ^15^. A low dose, short treatment paradigm did not alter the proliferation of human cortical progenitor cells or induce apoptosis in human cortical neurons ^3^ and orally administered branaplam was also reported to have no impact on neurogenesis in the CNS in juvenile mice, rats, and dogs ^16^. Our straight forward and efficient in vitro framework suggests clear neuropathogenic effects, including detection of elevated levels of neurofilament light chain, a clinically accepted biomarker for neurite degeneration. This effect was also observed in a more elaborate dog model upon branaplam administration over the course of several ^17^. This reflects the severe affection of the peripheral nervous system and specifically axons in HD patients that led to the termination of the VIBRANT-HD study. Multiple reasons may act synergistically. First, the pharmacokinetic profile of infants vs. adults may be different. Additionally, HD patients may be more susceptible to branaplam-induced neuropathy. Mutant HTT directly interacts with p53 and this interaction may contribute to neurodegeneration ^18, 19^. Hence, interfering with p53-dependent pathways through small molecules may be problematic in HD. Lastly, more subtle peripheral neuropathic effects, and especially motor neuropathy, may have been missed in infants with SMA, due to the disease phenotype of motor neuron degeneration versus potential degeneration of the motor axon due to a drug’s side effect. Potential protective effects of elevated SMN2 levels may have masked toxic changes induced by branaplam. To finally resolve this issue, more detailed investigations will be necessary and will require in-depth analyses and comparison of the clinical parameters of the branaplam trials in SMA infants (NCT02268552) and HD patients (NCT05111249).

On a molecular level, these side effects may arise due to mRNA-independent mechanisms of branaplam. Multiple drugs lead to peripheral neuropathy as a side effect, most prominently chemotherapeutics such as taxans and platin-containing substances. While taxans are thought to directly destabilize microtubules, platin-containing chemotherapeutics exhibit manifold cytotoxic effects ^20^. Importantly, oxaliplatin very frequently leads to peripheral neuropathy. It has been shown to induce nucleolar stress measured via translocation of NPM1 from the nucleolus to the nucleoplasm ^21–23^ and p53 stabilization ^21^, as observed in the present study. Oxaliplatin is thought to induce nucleolar stress by disrupting rRNA biogenesis ^21, 23^. Therefore, off-target effect mechanisms on non-coding RNAs may be involved and have to be considered, especially for drugs known to bind and interfere with RNA.

Due to the nucleic acid sequence and structure-dependent mechanisms, this illustrates the urgent need of involving human-based assays and the superior power of human in vitro models for testing on- and off-target effects of nucleic acid-targeting drugs.

## MATERIALS AND METHODS

### Study design

The objective of this study was to investigate the side effects of splicing modulator branaplam after the phase II study for the treatment of HD was stopped due to peripheral neuropathic symptoms in treated patients. The results were achieved from in vitro and in silico experiments. The sample size was based on previous experience in cell culture experiments. No data were excluded. In silico analysis was based on previously published RNA-seq datasets from human fibroblast (n=8 (n=4 Ctrl, n=4 HD) and human iPSCs-neurons (n=6 (n=3 Ctrl, n=3 HD), which were treated with DMSO or branaplam. First, we hypothesized mRNA-off target effects of branaplam. Therefore, we compared the neuropathy gene lists from Mayo Clinic Lab and MGZ Munich with our results. Our analysis did not reveal gene expression changes which are likely to explain the peripheral neuropathic side effects. Next, we hypothesized that branaplam treatment induces toxic side effects. Differentially gene expression analyses of the previously published RNA-seq data sets from fibroblasts revealed enrichment for the p53-regulated genes. ENCODE p53 ChIP-seq dataset was used to confirm the global p53 activation. Cell cycle analysis of human fibroblast was performed to in vitro confirm the p53-mediated cell cycle arrest upon branaplam treatment. To investigate p53-mediated neurotoxic effects upon branaplam treatment, we differentiated hiPSC-derived motor neurons from Ctrls and treated them with DMSO and branaplam. Nucleolar stress was quantified by immunofluorescence staining of NPM1 and the calculation of the ratio of NPM1 fluorescence intensity signal in the nucleoplasm and nucleolus. Neurotoxicity was analyzed by quantification of NfL in the supernatant and analyzing the neurite disintegration after immunofluorescence staining of beta-III-Tubulin in DMSO and branaplam-treated motor neurons. Protein expression of p53-target BB3 was quantified by western blot of whole motor neuron lysates treated with DMSO and branaplam.

### Differential gene expression analysis, overlaps and transcription factor binding site prediction

Differential gene expression was performed using DESeq2 (1.34.0)^24^ for fibroblasts (n=8 DMSO (n=4 Ctrl, n=4 HD); n=8 branaplam (n=4 Ctrl, n=4 HD)) and iPSC-neurons (n=6 DMSO (n=3 Ctrl, n=3 HD); n=6 branaplam (n=3 Ctrl, n=3 HD). A significant threshold was set to an adjusted P value of below 0.05. The overlaps of gene lists (branaplam targets, two neuropathy gene panel lists, differentially expressed genes in fibroblasts and neurons) was performed in Python and visualized as Venn diagrams using the matplotlib plugin.

Counts were transformed into RPKM (Reads Per Kilobase of transcript per Million reads mapped) values and the fold change of expression upon was calculated first for each individual and then averaged across the group (Ctrl or HD sample). Fold changes were plotted as cumulative distribution plots in Python using the ecdfplot function of the seaborn plugin. Significant differences were determined using 2 sample KS-test using scipy.stats. As a control fold change values of a shuffled list of genes with equal size as the tested condition was used. To predict enrichment of transcription factor binding sites (TFBS) in differentially expressed genes, up- and downregulated genes were plugged into the DAVID gene ontology server^25^ and investigated for enrichment of UCSC TFBS. As a background, we uploaded all genes expressed in our fibroblast dataset, defined as a mean RPKM > 0.5 of a gene across all samples.

### P53 ChIP-seq validation

For validation of p53 association to gene expression changes, a p53 ChIP-seq peak dataset (conservative IDR thresholded peaks) was downloaded from ENCODE (ENCFF488WQN). We considered a region of 5000bp upstream of the gene start (determined via gencode annotation version 26) as a region of interest for putative promoter regions using pybedtools intersect (u=True). Fold changes of the determined putative p53 targets were plotted as cumulative distribution plots in Python using the ecdfplot function of the seaborn plugin. Significant differences were determined using 2 sample KS-test using scipy.stats. As a control, values of a shuffled list of genes with equal size as the tested condition was used.

### Human fibroblast and iPSC-derived cell lines

The generation and use of these human cell lines were approved by the Institutional Review Board (Nr. 4120 and 259_17B: Generierung von humanen neuronalen Modellen bei neurodegenerativen Erkrankungen). We used 8 previously published fibroblast lines from Ctrl (UKERf33Q, UKERfB26, UKERf4CC, UKERf4L6) and individuals with HD (UKERf4Q4, UKERfOP5, UKERf59H, UKERf919) ^3^. We also used 5 previously published iPSC-derived motor neuron progenitor lines originating from 3 control individuals (UKERiff2-X-18, UKERiff2-SC-11, UKERi33Q-E-2, UKERi33q-E-6, CV-B) ^5^.

### Fibroblast culture

Fibroblasts were resuspended in fibroblast growth medium (FGM; 75% DMEM, 15% FCS, 2 mM l-glutamine, 100 µg/ml penicillin/streptomycin, 2 ng/ml fibroblast growth factor 2) and plated on polystyrene cell culture flasks. Medium was changed twice a week. To expand the fibroblasts, they were split by washing the cells once with PBS (w/o CaCl2, w/o MgCl2), followed by incubation with Trypsin at 37 °C for 20 min. The detached cells were collected in DMEM, transferred to a centrifugation tube and centrifuged for 51min at 3001×g at room temperature. Afterwards, the supernatant was removed, cells were resuspended in FGM and plated on a new polystyrene cell culture flask. For imaging-based experiments, fibroblasts were seeded on a 96-well glass bottom imaging plate (1000 cells per well). The cells were treated with three different concentrations of branaplam (treatment concentrations: 10 nM, 100 nM and 1000 nM) as well as the vehicle (DMSO). For flow cytometry, cells were seeded on a 10 cm dish (200,000 cells per dish) and treated with branaplam (10 nM, 100 nM and 1000 nM), the vehicle (DMSO) and paclitaxel for 72 hours. All treatments were performed 72 h, with complete media changes every 24 h.

### Motor neuron differentiation

Frozen stocks of motor neuron progenitor (MNP) cells were differentiated from iPSC as described in detail previously ^5^. Frozen MNP were thawed by adding 1 ml of warm DMEM/F12 + Glutamax to a semi-thawed vial. The cells were collected in a 15 ml Falcon tube, 5 ml of warm DMEM/F12 + Glutamax were added to dilute the DMSO in the freezing media and centrifuged at 200g for 3 min at room temperature. The supernatant was aspirated and resuspended in 1 ml of a basal media, further referred as N2B27 (DMEM/F12, 1xN2, 1x B27, 100 μM ascorbic acid, 1xPen/Strep) supplemented with 1.5 μM retinoic acid (RA) and 200 nM of Smoothed agonist (SAG) 10 μM Rock inhibitor (RI) and 2 ng/ml of brain-derived neurotrophic factor (BDNF), glial-derived neurotrophic factor (GDNF), and ciliary neurotrophic factor (CNTF), respectively. This combination of growth factors will be referred to as neurotrophic factors (NFs) further on. 35,715 cells/cm^2^ were seeded on Geltrex-coated 12-well plates and cultured with 1 ml media per well. Two and 4 days after seeding, media was exchanged completely and the cells were fed with N2B27 and 1.5μM RA, 200 nM SAG, 10 μM RI and NFs. From day 6 on, cells were cultured in N2B27 and 2 μM DAPT, 10 μM RI and NFs. On day 7 after seeding, an additional feeding step was performed with cold N2B27 and 2 μM DAPT, 10 μM RI and NFs plus Geltrex and 0.5 μg/ml laminin to enhance cell attachment. On day 9, the cells were washed 1x with DMEM/F12 + Glutmax and then fed further on with N2B27 and 10 μM RI, NFs and 0.5μg/ml laminin till the end of differentiation (day 14 after seeding). For imaging-based experiments, 6 days after seeding cells were dissociated with Accutase, and plated on Geltrex coated cover slips in 24-well plates (50,000 cells per well) or 96-well imaging plates (25,000 cells per well). Branaplam treatment (10 nM, 100 nM, 1000 nM) or vehicle (DMSO) was administered to the cells on day 9, 10 and 12 after seeding by exchanging the media completely.

### MSD^®^ Multi-Spot Assay System

Multiplex ELISA was performed using the Meso Scale Discovery^®^ system (MSD^®^; Rockville, MD, USA) according to the manufacturer’s recommendation. For this experiment the U-Plex^®^ Development Pack, 96-well 2-Assay SECTOR Plate kit (Cat.: K1522N) was used together with the R-PLEX Human Neurofilament L Antibody Set (Cat.: F217X) and recommended buffers, Diluent 11 (Cat.: R55BA) and Diluent 12 (Cat.: R50JA). For the analysis, cell culture supernatants were diluted 1:2 in Diluent 12. All steps were performed according to the manufacturer’s instructions. Provided plates were coated with Linker-coupled antibodies one day before the assay. The biotinylated antibody was combined with the assigned Linker (Linker 1 for NF-L) and adjusted to 6 ml with the provided Stop Solution. 50 µl of the coating solution was added to each well and the plate was subsequently incubated for 1 h at RT and afterwards washed three times with PBS 0.05 % Tween-20. The plate was stored at 4 °C over night. For the assay, provided calibrators were diluted in a 4-fold serial dilution using Diluent 12 to generate eight standards. Next, 25 µl of Diluent 12 was added to each well, followed by 25 µl of prepared calibrator standards or sample dilutions. All standards and samples were added in duplicates. The plate was incubated at RT with shaking for one hour. After the plate was washed three times as described above, 50 µl of detection Antibody Solution was added to each well. The 100X stock solution of the detection antibody was diluted in Diluent 11 shortly before. After an incubation for 1 h, the plate was again washed three times and 150 µl of provided MSD GOLD Read Buffer B was added to each well. The measurement was performed by MESO^®^ QuickPlex^®^ SQ 120MM (Cat.: Al1AA) and subsequent analysis was carried out using MSD^®^ Discovery Workbench^®^ Version 4.0. All measured concentrations used for further analysis were within the manufacturer’s recommended detection range.

### Immune fluorescence imaging

Cells were fixed in 4 % paraformaldehyde (PFA) for seven minutes at room temperature and subsequently washed 3x with PBS for 3 mins. A total of five different Ctrl lines were processed in parallel. The cells were permeabilized and blocked in 0.3 % Triton X-100 and 5 % donkey serum in PBS for 30 mins at room temperature. Afterwards, the cells were incubated with primary antibodies (Nucleophosmin: ab10530, Abcam, 1:250; beta-III-Tubulin: ab18207, Abcam, 1:200; TAU: sc-1995, Santa Cruz Biotechnology, 1:500; ISL1: 39.4D5-s, DSHB, 1:250) in 5% donkey serum in PBS at 4 °C overnight. After washing twice with PBS for 3 mins, incubation with secondary antibodies (donkey anti-mouse IgG (H+L) secondary antibody, Alexa Fluor 488 conjugate: Thermo Fisher Scientific, A-21202, 1:500; donkey anti-goat IgG (H+L) secondary antibody, Alexa Fluor 546 conjugate: Thermo Fisher Scientific, A-11056, 1:500; donkey anti-Rabbit IgG (H+L) Secondary Antibody, Alexa Fluor 647 conjugate: Thermo Fisher Scientific, A-31573, 1:500) for 1 h at room temperature was performed. Then cells were washed once with 0.1 % Triton X-100 and 5 % donkey serum in PBS, followed by the nuclei staining using 0.2 µg/ml DAPI in PBS. Cells were washed twice with PBS, followed by mounting the cover split on the object slide using Mowiol solution. Imaging was performed with a Zeiss Observer.Z1, including Apotome technology.

### NPM1 staining analysis

Images were taken using a Zeiss Observer.Z1 using Apotome with a 63x objective. For each condition and line, 20 fields of view were imaged using 29 slices (14 µm) Z-stacks. After Apotome correction, a maximum intensity projection was applied and the images were exported as tiff files.

Nuclei and neurons were identified using IdentifyPrimaryObjects with the DAPI and beta-III-Tubulin images (DAPI: global Otsu thresholding with 2 classes, smoothing scale: 0.5, correction factor: 2; beta-III-Tubulin: adaptive Otsu thresholding with 2 classes, smoothing scale: 1.3488, correction factor: 1.3) and nuclei that belong to neurons were determined (RelateObjects). The NPM1 image was masked for areas only belonging to neuronal nuclei (MaskImage), enhanced (EnhanceEdges, Sobel method) and a gaussian filter was applied (Sigma: 2.5). Next, primary objects (nucleoli) were determined using IdentifyPrimaryObjects (adaptive Otsu thresholding with 2 classes, smoothing scale: 1, correction factor: 1, do not allow filling holes in objects), merged into a single object (SplitOrMergeObjects) and holes within Objects are filled (FillObjects, maximum hole size: 50). The nucleoplasm area (non-nucleolar nucleus area) was identified using IdentifyTertiaryObjects. The MeasureObjectSizeShape and MeasureObjectIntensity were subsequently applied. A ratio of NPM1 fluorescent intensity in the nucleoplasm that is not the nucleolus (non-nucleolar) vs. the nucleolar area was calculated for each cell. The median for this ratio was calculated for each line and condition and used for statistical analysis.

### Neurite disintegration analysis

Images were taken using a Zeiss Observer.Z1 using Apotome with a 20x objective. For each condition and line, three to four fields of view were imaged using 50 slices (36,75 µm) Z-stacks. After Apotome correction, a maximum intensity projection was applied and the images were exported as a tiff file.

Quantification of neurite disintegration was performed on basis of the quantification of the microtubule depolymerization index ^14^. The analysis was performed in CellProfiler (4.1.3). The analysis is based on that destabilized microtubules such as in damaged neurites and neurite swellings occupy higher intensity values. In a first step, the intensity and standard deviation of intensity in control conditions (DMSO) within neurites is determined to set an intensity threshold (3 standard deviations above the mean intensity) to determine occupancy of pixels to the highest intensities in neurite occupying areas. This threshold, determined for each cell line separately, is then applied to all images and the area occupied by disintegrated neurite areas is divided by the total neurite area in an image and defined as the neurite disintegration index. Specifically, the original input image was subject to the EnhanceOrSupressFeatures function to enhance neurites (Feature type: Neurites; Enhancement method: Line structures; Feature size: 10). This was used to determine a binarized object of interest (neurite area) using the IdentifyPrimaryObjects function (Threshold strategy: Global; Threshold method: Otsu with two classes; Smoothing scale: 0; Threshold correction factor: 1.3; no filling of holes in objects) and merged into a single object (SplitOrMergeObjects). The fluorescent intensity and standard deviation thereof in this area was quantified (MeasureObjectIntensity). For each cell line, the results from the DMSO images were averaged separately to determine a cell line-specific intensity threshold (3 standard deviations above the mean intensity). In a second step, all images were processed from the start again in the same fashion as described before. A mask of the binary object of interest (neurite area) was applied to the original beta-III-Tubulin image. The cell line specific intensity threshold was applied to the masked image to obtain a binary image of disintegrated neurite areas (IdentifyPrimaryObject; Threshold strategy: Global; Threshold smoothing scale: 0) and merged into a single object per image (SplitOrMergeObjects). The area occupied by the total neurite area and the disintegrated area was determined (MeasureObjectSizeShape) and exported. A disintegrated neurite to total neurite area ratio was applied for each image (AreaShape_Area columns used) and averaged for each cell line.

### Western blot

For whole cell lysates, cells were lysed in Radio-immunoprecipitation buffer (RIPA: buffer ingredients: 50mM Tris-HCl (pH7.4), 1% NP-40, 0.5% Deoxycholic acid sodium salt, 0.1% Sodium dodecyl Sulfate (SDS), 150mM Sodium chloride, 2mM Ethylenediaminetetraacetic acid (EDTA), 50mM Sodium fluoride) followed by sonication with a Diagenode Bioruptor Pico (setting: 30s ON, 30s OFF, 5 cycles, high frequency). The protein concentration was estimated by bicinchoninic acid (BCA) assay and equal concentrations were applied. All immunoblots were run on 4–12% Bis-Tris gels with NuPAGE MOPS running buffer for 901min at 1501V. Proteins were transferred to a PVDF membrane with 10% methanol in NuPAGE transfer buffer at 10 V for 14 h at 4 °C. The membrane was then blocked for 1 h in 5% dry milk in TBS-T. Afterwards, primary antibodies (BBC3 (PUMA): ab9643, Abcam, 1:500; GAPDH: CB1001, Millipore, 1:5000) in 5% dry milk in TBS-T were incubated for 1 h at room temperature. Subsequently, the membrane was washed twice for 3 mins in TBS-T and then incubated with the secondary HRP-conjugated antibody (donkey anti-mouse IgG (H+L) secondary antibody, HRP, Thermo Fisher Scientific, SA1-100, 1:5000; Donkey anti-rabbit IgG (H+L) secondary antibody, HRP, Thermo Fisher Scientific, SA1-200, 1:5000) in 5% dry milk in TBS-T for 1 h at room temperature. The membrane was washed three times for 3 mins with TBS-T and incubated in the dark with ECL solution. The chemiluminescent signal was detected using a film and developed in the dark with various exposure times. Western blot signals were quantified densitometrically using Fiji. The signal was normalized to the corresponding signal of GAPDH.

### Cell cycle assay

For cell cycle analysis, human fibroblasts were cultured and treated as described before. For EdU incorporation, culture media was supplemented with 10 µM 5-Ethyl-21-deoxyuridine (EdU) for one hour at 37 °C. Cells were immediately dissociated after incubation as described before and resuspended in FC buffer (2 % FCS, 0.01 % sodium azide, 3 mM EDTA). The suspension was dispensed into 5 ml tubes (Sarstedt) at 5×10E5 cells per tube. Cells were fixed and permeabilized in 100 µl BD Fixation/Permeabilization Solution (BD Bioscience) for 10 min. Then 1 ml of BD Perm/Wash Buffer (BD Bioscience) supplemented with 0.1 % Triton X-100 was added to and cells were centrifuged at 1,700 rpm for 3 min. EdU labeling was performed with EdU Click FC ROTI®kit for Flow Cytometry (Carl Roth) according to the manufacturer protocol. EdU-labeled cells were stained with the click assay cocktail for 30 min at room temperature, protected from light. Subsequently, 1 ml of BD Perm/Wash Buffer (BD Bioscience) supplemented with 0.1 % Triton X-100 was added and cells were centrifuged at 1,700 rpm for 3 min. For DNA staining, cells were resuspended in 1 ml FC buffer supplemented with 2 µg/ml DAPI. After 30 min incubation at RT, protected from light Flow cytometry was performed with Cytoflex S. Sample flow rate was adjusted to a maximum count of 400 events per second for singlet counting. Analysis was performed with CytExpert Software. Cell cycle phases were gaited from gaited single nuclei events, according to their DNA content (2n=G0/G1, 4n=G2/M) or EdU positive events (S-Phase) (Supplementary Fig. 2).

### Statistical analysis

GraphPad Prism 9 was used to visualize data and calculate statistics for pairwise and grouped analyses (RPKM analysis of specific genes, FACS quantification, densitometric quantification of western blot, image data analyses). DMSO samples and their respective treated samples were considered as paired. When comparing two conditions, Mann–Whitney–U test was used. When comparing multiple groups (e.g. different branaplam concentrations), Friedman test was used for non-normally distributed data with Dunn’s post hoc test, respectively, to identify differences between individual groups. For grouped analyses (e.g. DMSO vs. branaplam in Ctrl vs. HD), paired two-way ANOVA was used. The statistical test and exact P value used for calculating the significance of each graph is indicated in the figure legend. A P value1≤10.05 was considered as significant.

## Acknowledgments

We thank Naime Zagha, M.Sc., Sonja Ploetz, and Michaela Farrell for excellent technical support. Data analysis was performed at the servers hosted by the “Data Integration Center” (DIC) and accommodated by Dr. Wolfgang Krebs and Dr. Pooja Gupta within the “Core Unit for Bioinformatics, Data Integration and Analysis” (CUBiDA), Universitätsklinikum Erlangen, Erlangen, Germany.

## Author contribution Conceptualization

FK, TB, JW, BW Formal analysis: FK

Investigation: FK, TB, SR, LK Resources: MR, JW, BW

Funding acquisition: FK, JW, BW, MR Supervision JW, BW

Writing – original draft: F.K. and T.B. made figures and wrote the original draft of the manuscript.

Writing – review & editing: FK, TB, MR., JW, BW

## Data and materials availability

All data to generate graphs are provided in the source data file. Counts tables of fibroblasts and iPSC-neurons treated without and with branaplam and branaplam’s targets from our previous study were used and the respective counts are publicly available in the source data of that paper ^3^. Gene lists associated with hereditary neuropathy were acquired from gene panel diagnostics from the Mayo Clinic Laboratories (’Peripheral Neuropathy Expanded Panel’; list from 2019; https://www.mayocliniclabs.com/) and the MGZ Munich (’Neuropathy/motor neuropathy comprehensive panel’, list available in November 2022, https://www.mgz-muenchen.com/). The p53 ChIP-seq dataset (ENCFF488WQN) was downloaded from the ENCODE project ^8, 9^.

Additional information, and materials that support the findings of this study are available from the corresponding author upon reasonable request.

## Supplementary material

**Fig S1.**
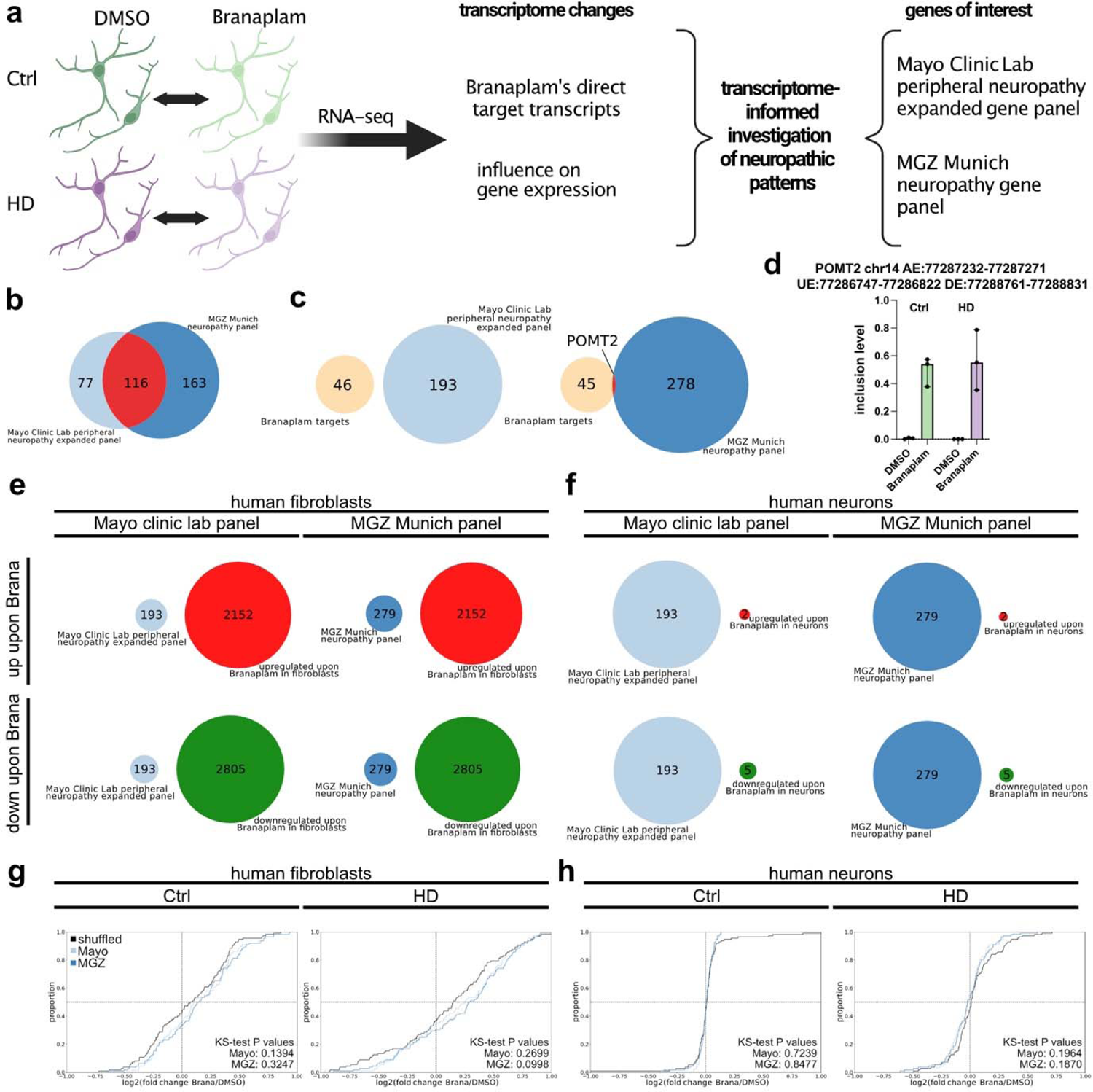
Branaplam does not directly target hereditary neuropathy associated genes. (A) Paradigm of analysis strategy. Published branaplam target genes and fibroblast and iPSC-neuron RNA-seq data from Ctrl (greens) and HD (purples) with DMSO or 72h 10 nM branaplam treatment (opaques) was used ^3^ and compared to two genes in two diagnostic panels for hereditary neuropathies. (B) Venn diagram showing overlap of genes (red) of the two neuropathy gene panels (light blue: Mayo Clinic Lab; blue: MGZ Munich). (C) Venn diagram showing overlap (red) of previously determined branaplam targets (orange)^3^ and gene list from Mayo Clinic Lab (light blue, left) or MGZ Munich (blue, right). (D) Bar graph with inclusion levels of branaplam-targeting POMT2 exon in iPSC-neurons of Ctrl (greens) and HD (purples) with DMSO or branaplam (opaques) treatment. Top depicts exact location of event in GRCh38. AE: alternative exon, UE: upstream exon, DE: downstream exon. Data shown as median and interquartile range. (E) Venn Diagrams of overlap of differentially upregulated (red) or downregulated (green) genes (DESeq2 adjusted P value < 0.05) upon 72h 10 nM branaplam treatment in fibroblasts with Mayo Clinic Lab (light blue) or MGZ Munich (blue) neuropathy gene panel list. (F) Venn Diagrams of overlap of differentially upregulated (red) or downregulated (green) genes (DESeq2 adjusted P value < 0.05) upon 72h 10 nM branplam treatment in iPSC-neurons with Mayo Clinic Lab (light blue) or MGZ Munich (blue) neuropathy gene panel list. (G) Cumulative distribution plot of Ctrl (left) or HD (right) fibroblast log2(fold changes) of branaplam vs. DMSO of genes in the Mayo Clinic Lab (light blue) or MGZ Munich (blue) list and randomly shuffled background of equal size (black). Significance calculated with 2 sample Kolmogorov-Smirnov (KS) test compared to shuffled background. (H) Cumulative distribution plot of Ctrl (left) or HD (right) iPSC-neurons log2(fold changes) of branaplam vs. DMSO of genes in the Mayo Clinic Lab (light blue) or MGZ Munich (blue) list and randomly shuffled background of equal size (black). Significance calculated with 2 sample Kolmogorov-Smirnov (KS) test compared to shuffled background.

**Fig. S2:**
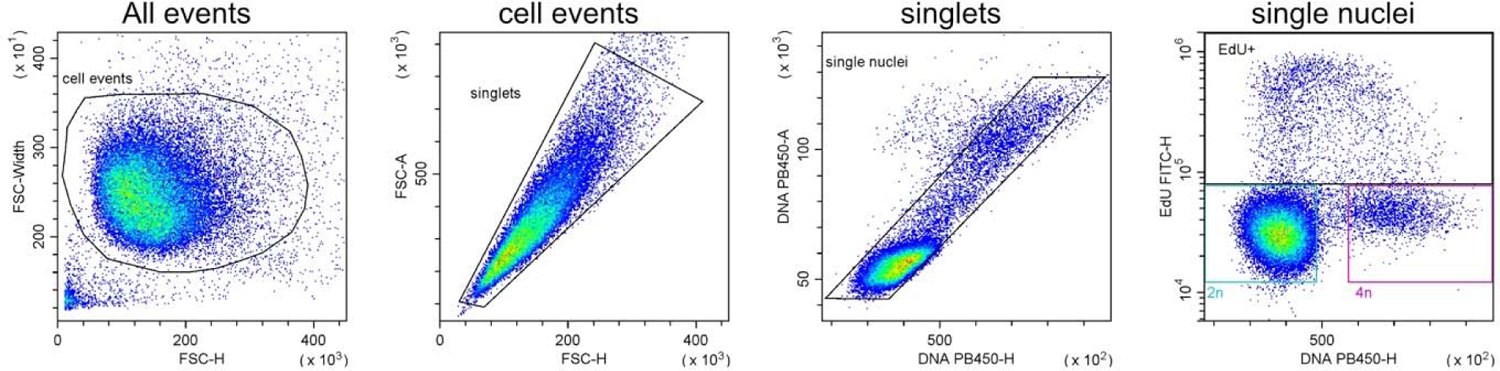
FACS analyses for cell cycle assay

Gating strategy for cell cycle assay of human fibroblasts.

